# Cortical encoding during word reading in bilinguals: parallel interdigitated distributed networks support convergent linguistic functioning

**DOI:** 10.1101/2021.11.25.470068

**Authors:** Shujie Geng, Wanwan Guo, Kunyu Xu, Tianye Jia, Wei Zhou, Colin Blakemore, Li-Hai Tan, Miao Cao, Jianfeng Feng

## Abstract

Word reading includes a series of cognitive processes that convert low-level visual characteristics to neural representations. However, the consistency of the neural mechanisms for processing these cognitive components across different writing systems in bilinguals remains inconclusive. Here, we explored this question by employing representational similarity analysis with a semantic access task involving Chinese words, English words and Chinese pinyin. Distinct anatomical distribution patterns were detected for each type of brain representation across ideographic and alphabetic languages, resulting in 100% classification accuracy. Meanwhile, convergent cognitive components processing was found in the core language-related regions in left hemisphere, including the inferior frontal gyrus, temporal pole, superior and middle temporal gyrus, precentral gyrus and supplementary motor areas. Broadly, our findings indicated that the neural basis for word recognition of different writing systems in bilinguals was divergent in anatomical locations of neural representations for specialized processing but convergent in cognitive domains, which supported and enriched the assimilation-accommodation hypothesis.

**Teaser:** Cortical encoding linguistic processing across languages was support by parallel interdigitated distributed networks but convergent in cognitive domains.

## Introduction

Reading, a highly complex and advanced learned ability, is vital for new knowledge acquisition and problem solving. Three language-processing components, orthographic, phonological and semantic representations, were found to be processed and integrated in brain language networks during reading to integrate information from visual properties and pronunciations of printed words and access semantics (J. Dong et al., 2021; Fischer-Baum, Bruggemann, Gallego, Li, & Tamez, 2017; Price, 2012; Rayner, Foorman, Perfetti, Pesetsky, & Seidenberg, 2001). However, the key question of whether neural populations supporting these processing components in different writing systems in bilingual individuals are convergent is still in debated.

Specifically, English is an alphabetic language whereas Chinese is an ideographic language (Yu & Reichle, 2017). These two writing systems vary significantly in visual-spatial properties, orthographic rules (i.e., the regularity of mapping from graphemes to phonemes) and semantic access (Perfetti, Liu, & Tan, 2005; Reichle & Yu, 2018; J. Zhao et al., 2014). Although previous neuroimaging studies suggested a similar pattern of cortical areas associated with Chinese and English reading, including the left inferior frontal gyrus, left superior and middle temporal gyrus, left angular gyrus and left middle fusiform gyrus (Bolger, Perfetti, & Schneider, 2005; Cao, Brennan, & Booth, 2015; Jobard, Crivello, & Tzourio-Mazoyer, 2003; Price, 2012; Rueckl et al., 2015), Chinese and English employ differential neural mechanisms that could be linked with linguistic attributes at the word reading level. English reading relies more heavily on the orthography-to-phonology mapping pathway and phonological awareness than Chinese reading does (Ruan, Georgiou, Song, Li, & Shu, 2018). In contrast, Chinese word reading requires a person to map the visual forms onto a whole syllable and depends more on the semantics-mediated pathway and morphological awareness than English reading does (C. S. Chen, Xue, Mei, Chen, & Dong, 2009; Ruan et al., 2018; Tan, Laird, Li, & Fox, 2005). Hence, specific activations for English reading were found in brain areas for phonological processing such as the left precentral gyrus, dorsal inferior frontal gyrus, supramarginal gyrus and temporoparietal cortex (Bolger et al., 2005; Cao et al., 2015; Cummine et al., 2013; Jobard et al., 2003; Price, 2012; Tan et al., 2005). Moreover, Chinese reading elicits more activations in the bilateral fusiform gyrus, right middle occipital gyrus, left ventral inferior frontal gyrus, and lateral temporal gyrus; these areas provide orthographic and semantic processing (Bolger et al., 2005; J. Dong et al., 2021). Explorations conducted in bilingual populations also confirmed the intermingled neural populations responding to Chinese and English word reading (Rueckl et al., 2015; Xu, Baldauf, Chang, Desimone, & Tan, 2017). The assimilation-accommodation hypothesis proposed by Perfetti and colleagues indicated that the human brain not only utilizes neural networks supporting native language to complete second language reading (known as assimilation), but also recruits new neural networks to adapt unique linguistic features of the second language (known as accommodation) (Perfetti et al., 2007). In addition, language acquisition order and learning experience would in turn affect the brain language-processing mechanisms for the first language. For example, after long-term learning of Chinese in adulthood, English native speakers were shown to exhibit greater activation bilaterally in the fusiform gyrus during English reading (Mei et al., 2015). Notably, Chinese pinyin, as a phonetic symbol system for Chinese characters, shares similar alphabetic orthography-phonology mapping properties with English words but uses the same phonology and semantic lexicons as Chinese words (Ding et al., 2018). Overuse of Chinese pinyin as input negatively affected behavioral performance and neurodevelopment for Chinese reading in children (Li et al., 2020; Tan, Xu, Chang, & Siok, 2013). In this sense, the underlying neural mechanisms for Chinese pinyin processing might be different from both Chinese words and English words.

However, previous studies of this issue were limited to depictions of the overlapping and respective activation maps as well as related features (e.g., lateralization) between English and Chinese reading but were insufficient to explain whether the underlying neural mechanisms related to linguistic properties (i.e., orthographic, phonological and semantic representations) were consistent during language processing, especially for bilinguals (Bolger et al., 2005; Tan et al., 2003). To date, it is still largely unknown which cortical regions involved in each linguistic property for different writing systems, whether such neural basis accommodates linguistic features of writing systems and whether this basis is associated with behavioral performance.

Currently, beyond activation analysis, applications of representational similarity analysis (RSA) focus on explanations of brain activities on a certain theoretical model or property (Fischer-Baum et al., 2017; Kriegeskorte, Mur, & Bandettini, 2008; Mur, Bandettini, & Kriegeskorte, 2009; Wang et al., 2018; L. Zhao et al., 2017). Notably, algorithms in natural language processing have allowed us to establish quantitative measurements for complex cognitive linguistic models. These methods enable us to depict brain loads corresponding to different linguistic properties for Chinese and English reading. Here, we recruited a group of Chinese-English bilingual individuals who had received a unified education of Chinese pinyin at the elementary level in mainland China and passed College English Test Band 4. Participants performed a semantic access task for Chinese words, English words and Chinese pinyin during fMRI scanning (Figure 1A). The RSA method was used to quantify neural representations (i.e., involvement of specific language-processing components in language-related brain regions) of logo-grapheme, phonology, and semantic components for three writing systems. We hypothesized that 1) specialized neural representation patterns across writing systems exist, which accommodate to their own linguistic features 2) the divergent neural representation patterns are assimilated in the cognitive domains across writing systems.

**Figure 1.**
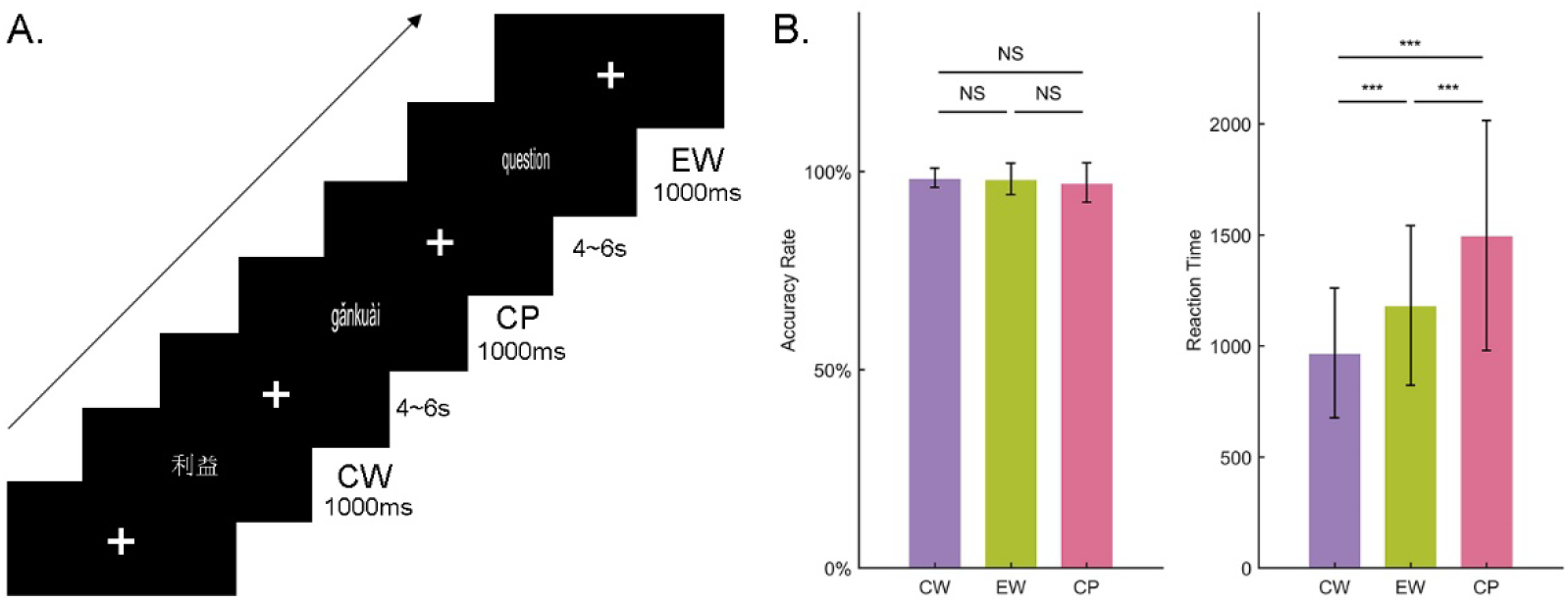
Experimental design and task performance. A. Event-related design was utilized in the current study. Visual stimuli of Chinese word, Chinese pinyin and English word reading tasks were presented for 1 s in randomized order with an inter-stimulus interval of 4~6s. Pertinently, responses were required to identify language types after semantic access. B. Accuracy rate and reaction time for Chinese word, Chinese pinyin and English word reading tasks. Reaction time was counted from the onset of visual stimuli to the button-press response. One-way ANOVA was conducted to test significant differences among language types. We found no significant differences among accuracy rates due to the ceiling effect but significant differences among reaction times. Post hoc analysis (Bonferroni corrected, p < 0.05) revealed the longest reaction time for accessing the semantics of Chinese pinyin and the shortest time for accessing the semantics of Chinese words. Group-level average reaction time indicated that subjects with full cooperation responded after semantic access. Abbreviations: CW, Chinese words; EW, English words; CP, Chinese pinyin. *** p < 0.001; ** p <0.01

## Results

First, we reported behavioral performances and group-level brain activation maps to depict task responses of all subjects (n = 51). To describe the underlying brain representations of different language components, representational similarity analysis (RSA) was utilized to determine the brain involvement of three components, including logo-grapheme, phonology and semantics, during the reading of Chinese word, English word and Chinese pinyin. We employed support vector regression (SVR) to reversely predict the condition category via a leave-one-out schematic. Finally, efforts were made to calculate the correlation between brain loads of cognitive components and behavioral performance (i.e., reaction time) for each condition.

### Behavioral performance and brain activations across three writing systems

The accuracy and reaction time results for the subjects are shown in Figure 1B. Group mean accuracy of Chinese word, English word and Chinese pinyin reading were 98.4% (87.5% - 100%, standard deviation (SD) = 2.4%), 98.1% (80% - 100%, SD = 4.0%), and 97.16% (80% - 100%, SD = 5.0%), respectively, and there was no significant difference across conditions (*F (2, 86)* = 1.20, *p* = 0.31). The group mean reaction time for Chinese word, English word and Chinese pinyin reading were 968.1ms (574.1 ms - 1769.8 ms, SD = 292.5ms), 1182 ms (681.9 ms - 1933.3 ms, SD = 359.4 ms), 1497.3 ms (718.1 ms - 2744 ms, SD = 518.3 ms), respectively. Significant differences across conditions were revealed by one-way ANOVA (*F (2, 86)* = 19.3, *p* < 0.01) and post-hoc two sample t-test (*p*s < 0.01, Bonferroni correction) with ascending reaction time for reading Chinese word, English word and Chinese pinyin, indicating the successful semantic access of all subjects when pressing a button.

Traditional brain activation maps at the group-level were calculated for each condition (Figure 2, *p* < 0.05 with false discovery rate (FDR) correction, cluster size > 10 voxels). Brain activation maps showed similar patterns across three language systems (Chinese word vs. Chinese pinyin, *r* = 0.90, *p* < 0.001; Chinese word vs. English word, *r* = 0.93, *p* < 0.001; and Chinese pinyin vs. English word, *r* = 0.97, *p* < 0.001). The middle temporal gyrus, fusiform areas, middle occipital gyrus, middle frontal gyrus, superior temporal gyrus, and inferior frontal gyrus were consistently activated under all conditions, which were in line with previous studies.

**Figure 2.**
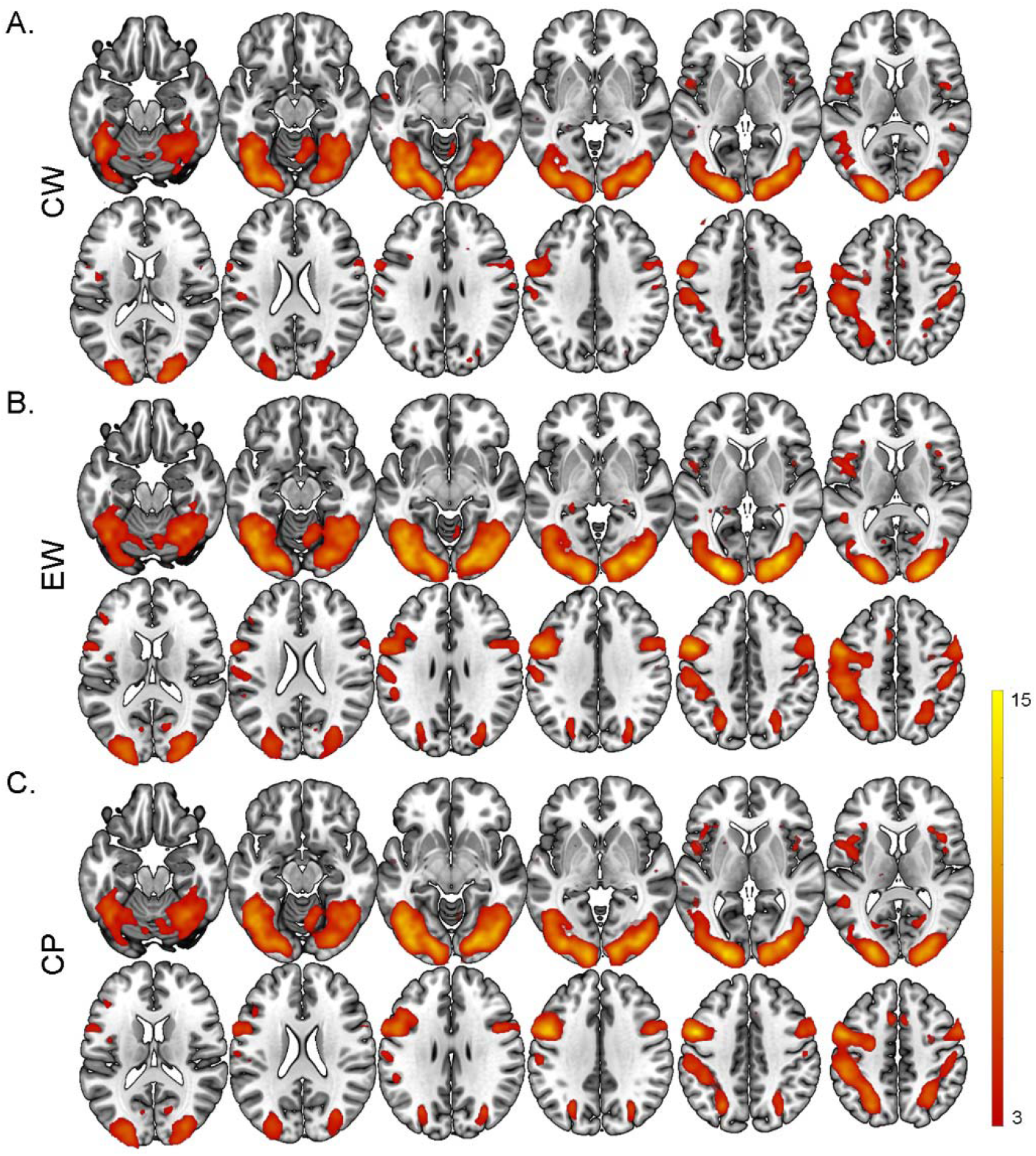
Activation analysis for Chinese word, Chinese pinyin and English word reading tasks. Second-level general linear models (GLMs) were performed separately to obtain activation maps for the recognition of A. Chinese words, B. Chinese pinyin and C. English words. Brighter colors indicate higher t values. (Voxel-wise p < 0.05, FDR corrected p < 0.05 and cluster size > 10). Abbreviations: CW, Chinese words; EW, English words; CP, Chinese pinyin.

### Distinguished representation patterns for language-processing components across writing systems

For RSA, firstly, we calculated the significant representation maps for all language-processing components across writing systems for each subject within a predefined language-related mask with the significance threshold set as *p* < 0.05 with cluster size > 10 voxels. The brain loads were then calculated as the sum of the representation values of all significant voxels. A classification analysis based on leave-one-subject-out cross validation and fold-specific mask was conducted to explore whether brain loads of separate language components were able to predict the input writing systems. We found that the category of ideographic and alphabetic writing system could be 100% correctly classified with the brain loads of all language components as features (accuracy: 100% between Chinese word and English word, 100% between Chinese word and Chinese pinyin, 96.1% between English word and Chinese pinyin, Figure 3A). Similar results were reproduced though employing the brain loads of each single component as features (accuracy based on logo-grapheme brain loads: 100% between Chinese word and English word, 100% between Chinese word and Chinese pinyin, 82.4% between English word and Chinese pinyin; accuracy based on phonology brain loads: 100% between Chinese word and English word, 100% between Chinese word and Chinese pinyin, 96.1% between English word and Chinese pinyin; accuracy based on semantic brain loads: 100% between Chinese word and English word, 100% between Chinese word and Chinese pinyin, 100% between English word and Chinese pinyin, Figure 3A). Furthermore, in recursive feature elimination (RFE) scheme based-on leave-one-sample-out cross validation and unified group-level mask, importance rank of features was identified. The brain areas that contributed significantly within the unified group-level mask are shown in Figure 3B, with the brain activity elicited by logo-grapheme processing in the left middle frontal cortices contributed the most.

**Figure 3.**
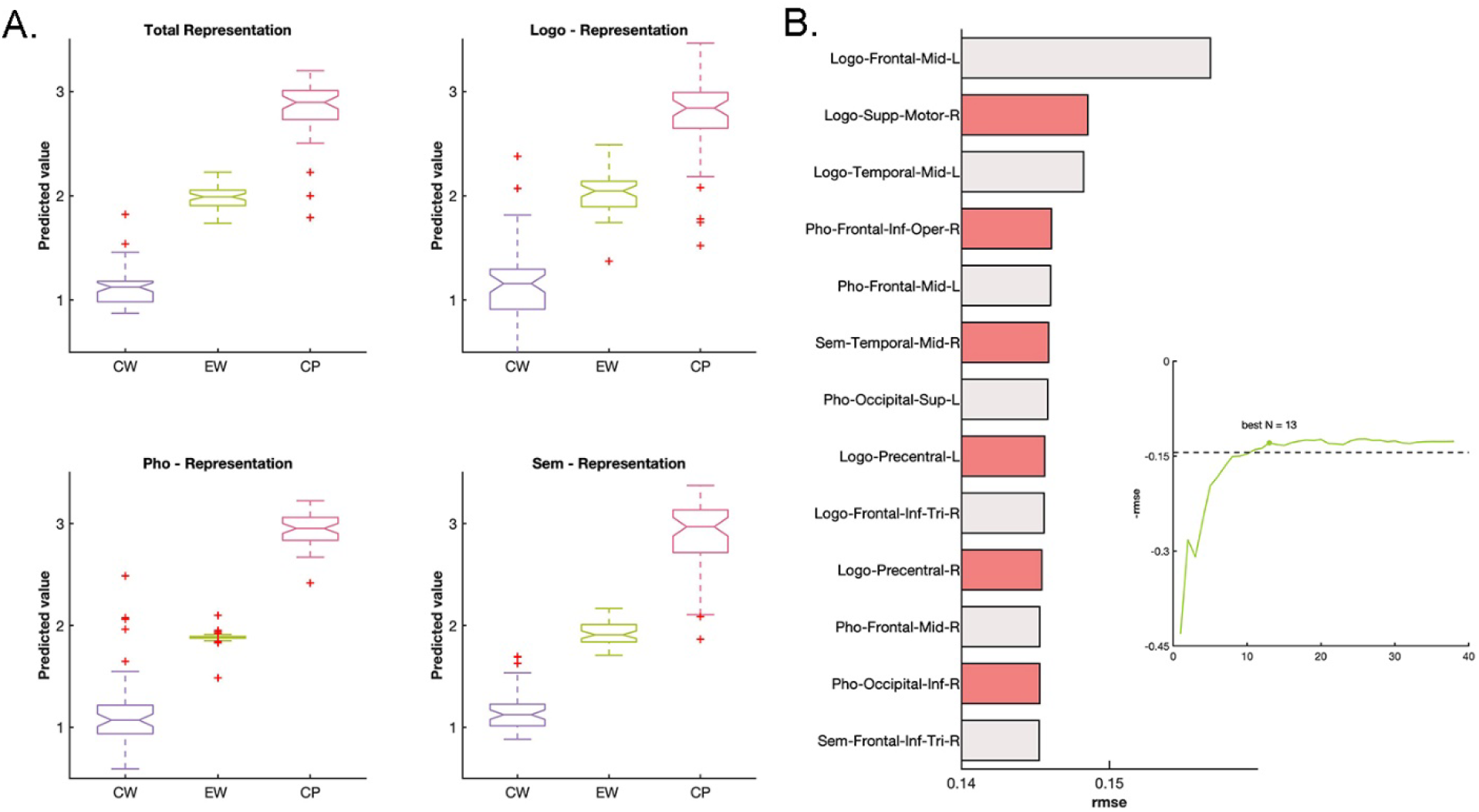
Separated brain activity patterns of language-processing components across language types revealed by the model SVR model. A. shows good predictions of language types regardless of the brain activity elicited by language-processing components in 30 ROIs as input features. B. Single features were recursively excluded to test its contribution to the model with total representations. A negative rMSE indicates the best number of features and ranks of features by contribution. Abbreviations: CW, Chinese words; EW, English words; CP, Chinese pinyin; Logo, logo-grapheme; Pho, phonology; Sem, semantics.

For more details, we showed the brain representation maps of each language system for every cognitive component of the group level in Figure 4A. The separated pattern across languages for all three components were detected throughout all cortical areas within the language mask. Similar anatomically separated patterns of linguistic components were also found at individual level. Interactions of brain loads between language cognitive components and writing systems were shown in Figure 4B. Chinese words reading showed equivalent brain loads for the three language components while Chinese pinyin and English words reading showed strikingly reversed brain loads in phonology. Note that for validation, these distinguished representation patterns were stable when the number of voxels was used as indexes to calculate brain loads.

**Figure 4.**
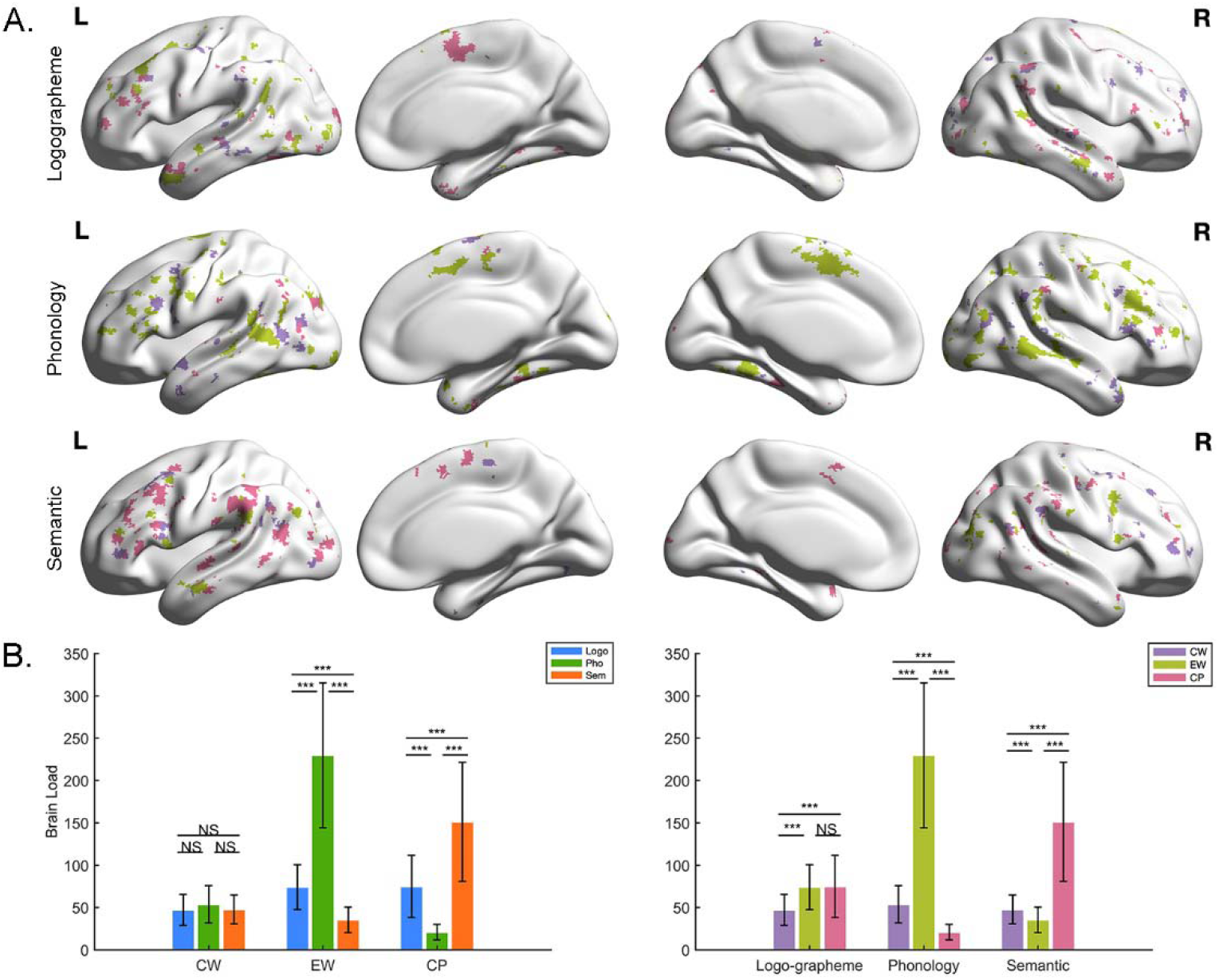
Brain loads of three language types. A. Brain involvement maps of 3 writing systems for each linguistic component. Panel. A shared figure legend with right part of panel. B. Purple indicates brain load elicited by Chinese, pistachio green represents English and magenta denotes Chinese pinyin. B. Interactions of brain loads between language types and linguistic components. Two-way repeated ANOVA was performed, and significant interactions were found between language-processing components and language types. Post hoc analysis shows differentiated patterns of brain activity elicited by language-processing components across language types: even patterns in Chinese words, V patterns in Chinese pinyin words and reversed V patterns in English words.

### Convergent cognitive domains for language-processing components across writing systems

From the perspective cognitive domain, we showed the brain representation maps of each language-processing components for each language system of the group level in Figure 5A. We found that most brain regions were involved in two or three language-processing components, indicating cognitive manifestation of specific component processing as well as transformations. Specifically, the cognitive manifestations in a majority of language-related regions were universal for the first and second languages, including the opercular parts of the left inferior frontal gyrus which were consistently involved in brain activity pertaining to phonological and semantic processing; the left precentral gyrus, left superior temporal gyrus, left middle temporal gyrus and left temporal pole which were consistently involved in brain processing of logo-grapheme, phonology and semantics (Figure 5B). Meanwhile, some regions were partially consistent in cognitive component involvement between the first and second language, which might be due to different demands of linguistic features (Figure 5C). The left middle frontal gyrus, left angular gyrus and left inferior parietal gyrus showed activity during logo-grapheme and semantic processing in Chinese reading but logo-grapheme and phonological processing in English; the triangular parts of the left inferior gyrus showed activity during phonology and semantics manifestations in Chinese reading but phonological, logo-grapheme and semantic processing in English reading. The left supramarginal gyrus demonstrated activity during phonology and logo-grapheme manifestations in Chinese reading but phonological and semantic processing in English reading.

**Figure 5.**
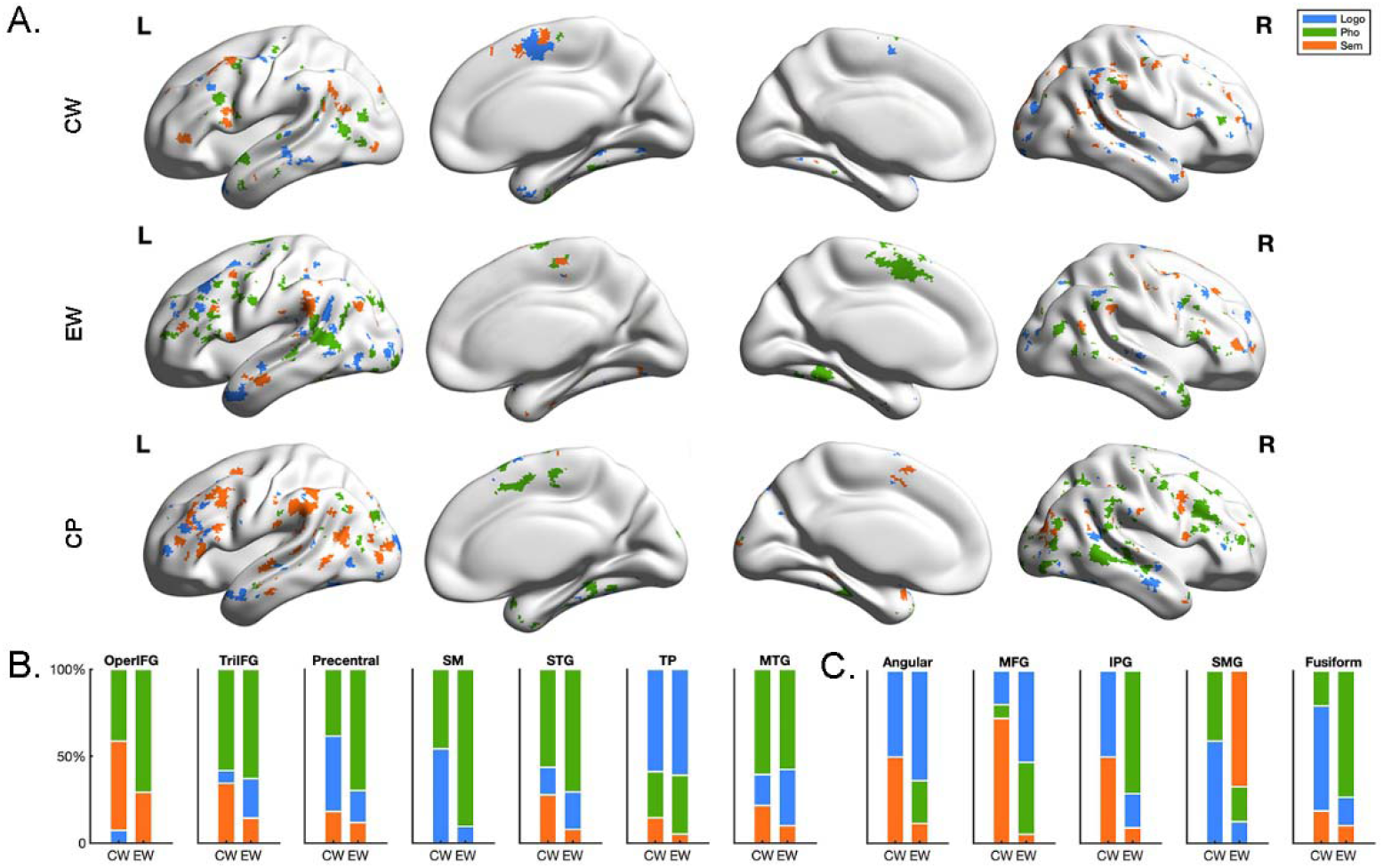
Brain loads of three language-processing components. A. Brain involvement maps of 3 language-processing components in each language type. B and C. The percentage of involvement of each component for language-related regions in Chinese word and English word reading. B indicates brain regions showed similar linguistic components manifestation between Chinese and English while C indicates brain regions with discrepancy manifestations.Blue indicates brain activity for logo-grapheme, green represents brain activity for phonology and orange denotes brain activity for semantics. Abbreviations: CW, Chinese words; EW, English words; CP, Chinese pinyin; Logo, logo-grapheme; Pho, phonology; Sem, semantics.

The overlapping areas of neural representations for all components between any two writing systems were also calculated (Figure 6 and Table S1). Shared brain involvement in core regions across writing systems were found. For logo-grapheme processing, overlapping brain representations were found in the right fusiform, right superior temporal cortex, left middle frontal cortex and right inferior frontal cortex for Chinese words and English words reading; the right superior temporal cortex, right inferior frontal gyrus (rIFG), right inferior parietal cortex and right precentral gyrus for Chinese words and Chinese pinyin reading; the left middle temporal cortex, left inferior occipital cortex, right superior temporal cortex for Chinese pinyin and English words reading. For phonological processing, overlapping brain activity were found in the right middle temporal cortex, right middle occipital cortex, left middle temporal cortex, left superior temporal cortex, left superior occipital cortex and right dorsal inferior frontal cortex for Chinese words and English words reading; the right fusiform, left superior occipital cortex, right supramarginal gyrus and right precentral gyrus for Chinese pinyin and English words reading. No overlap in brain activity was found between Chinese words and Chinese pinyin reading. For semantic processing, overlapping brain activity was found in the left precentral gyrus for Chinese words and English words reading; the left ventral inferior frontal gyrus, left middle temporal cortex, left angular cortex and right middle temporal cortex for Chinese words and Chinese pinyin reading; the left ventral inferior frontal gyrus, left supramarginal gyrus for Chinese pinyin and English words reading. Notably, only the triangular parts of the left inferior frontal gyrus corresponded to the semantic loads for all three writing systems, including Chinese words, English words and Chinese pinyin reading, while the overlaps failed to reach the threshold of cluster size larger than 10 voxels.

**Figure 6.**
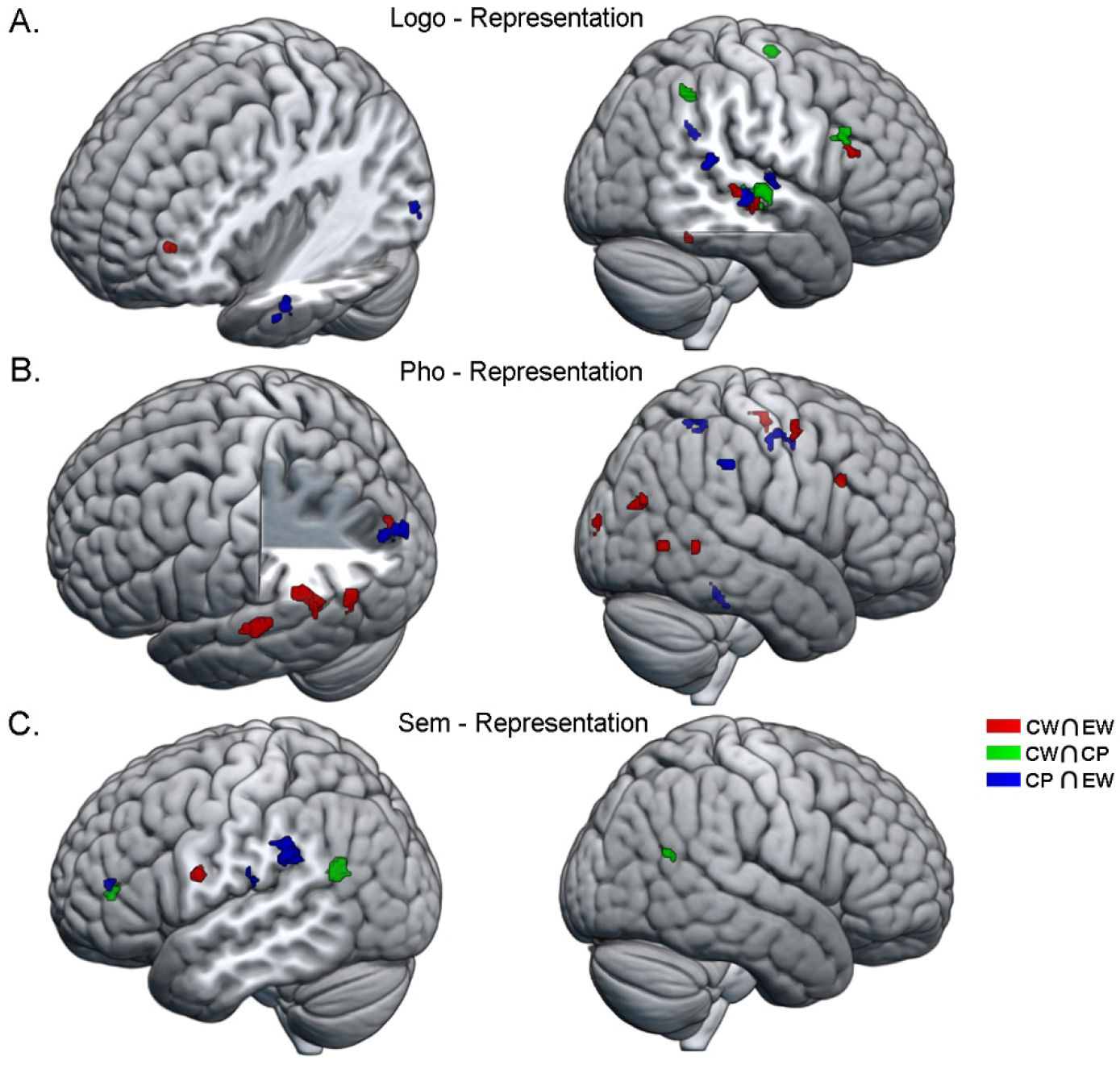
Overlap of brain involvement among language types in each language-processing component. Red represents overlapping brain involvement between Chinese word and English word reading; green signifies overlapping brain activity between Chinese word and Chinese pinyin reading; and blue indicates an overlap in brain activity between Chinese pinyin and English word reading. A. indicates brain involvement for logo-grapheme processing, B. depicts brain involvement for phonology and C. shows brain involvement for semantics. Abbreviations: CW, Chinese words; EW, English words; CP, Chinese pinyin; Logo, logo-grapheme; Pho, phonology; Sem, semantics.

Finally, for association analysis between brain and behavioral performances, we found significant partial correlations between the reaction time for Chinese word reading and logo-grapheme loads (*r* = 0.454, *p* < 0.05, Bonferroni corrected), Chinese pinyin reading and semantic loads (*r* = 0.528, *p* < 0.01, Bonferroni corrected), as shown in Figure 7. We also detected marginally significant correlations between reaction time for English word reading and phonological loads (*r* = −0.309, *p* = 0.047, uncorrected), Chinese pinyin reading and logo-grapheme loads (*r* = 0.303, *p* = 0.051, uncorrected). Pearson’s correlation analysis was also conducted between the total brain load and reaction time for every language type, but no significant correlations was found.

**Figure 7.**
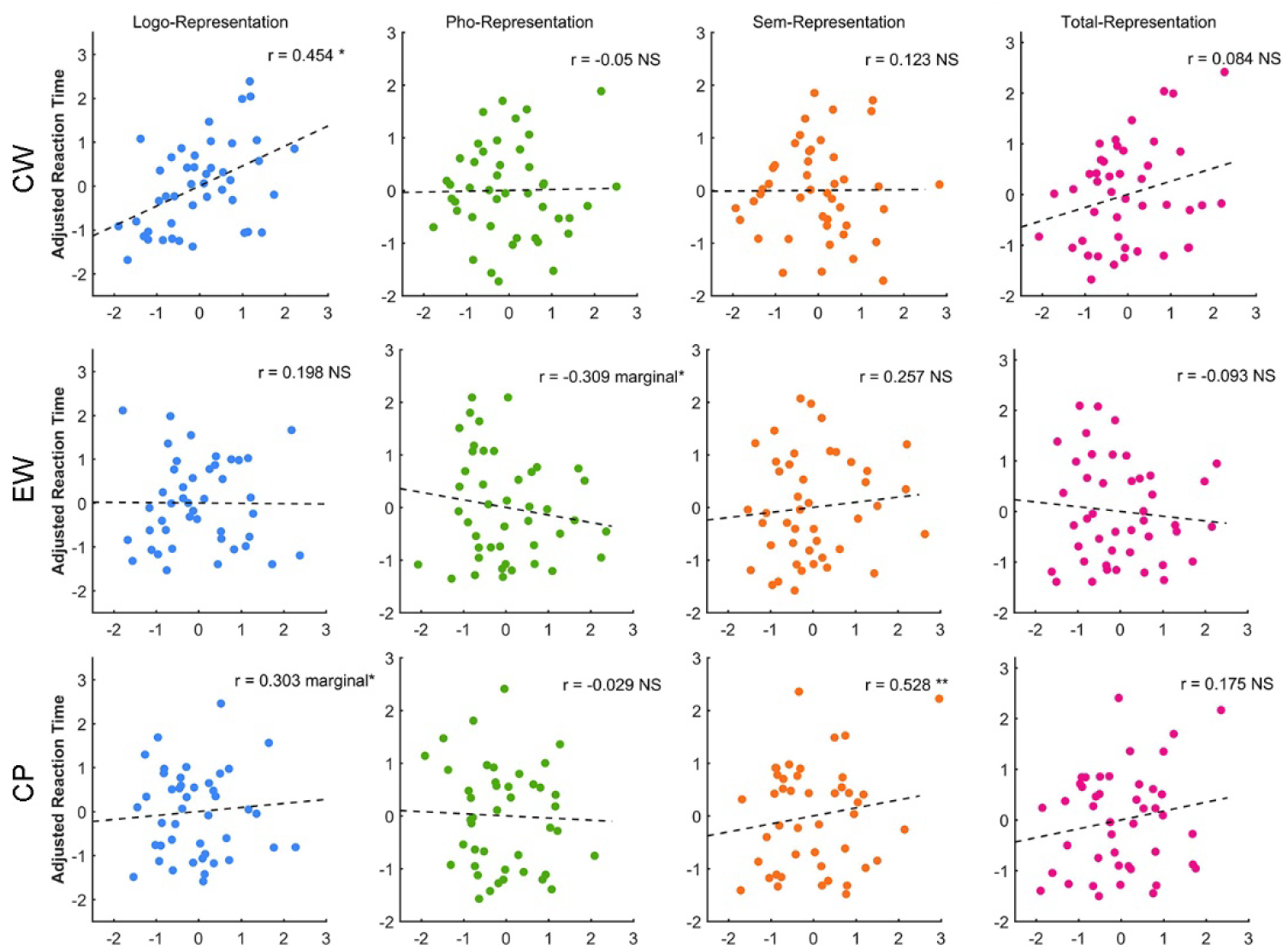
The association between brain activity loads with reaction times across three language-processing components/language types. Partial correlation was applied to calculate the relationship between brain activity elicited by language-processing components and reaction times for each language type. Abbreviations: CW, Chinese words; EW, English words; CP, Chinese pinyin; Logo, logo-grapheme; Pho, phonology; Sem, semantics.

## Discussion

In the current study, cortical encoding of three language cognitive components in Chinese-English bilingual individuals was investigated across Chinese word, English word, and Chinese pinyin reading tasks. In line with previous studies supporting the assimilation-accommodation hypothesis in bilingual individuals, we found both segregations and integrations between anatomical distribution and language cognitive components. Firstly, distinct spatial distribution patterns were detected for each type of brain representation across the three writing systems, resulting in 100% classification accuracy for the visual input category of the writing system. Language-specific differences were detected in the middle frontal gyrus, inferior parietal gyrus, angular gyrus, supramarginal gyrus, and fusiform in the left hemisphere. These differences might be due to the differences in linguistic properties. Meanwhile, convergence in cognitive components processing was found in other core language-related regions, including the opercular parts of the inferior frontal gyrus, the triangular parts of the inferior frontal gyrus, temporal pole, superior and middle temporal gyrus, precentral gyrus and supplementary motor areas. As a result of precision searchlight RSA, we depicted linguistic-feature specific divergence and convergence of neural populations responding to different writing systems. Cognitive fractionation of universal language system corresponded to the ‘assimilation’ theory while the specialization of anatomical representations for different systems, from more fine-grained perspective, provided evidence to ‘accommodation’ hypothesis.

We found that language types could be correctly classified with brain activity corresponding to all components as well as to every specific component as features, which indicates that not only different language components but also the same component across different language systems would entail distinguished neural populations which were adapted to their own linguistic features. Clinical findings were supportive for this result, that brain injuries may selectively impair the function of only one language in bilingual individuals; additionally, after operation therapy or brain stimulation, recovery of only one language may occur in bilingual language-impaired patients (Aladdin, Snyder, & Ahmed, 2008; Fabbro, 2001; Gomeztortosa, Martin, Gaviria, Charbel, & Ausman, 1995; Lucas, McKhann, & Ojemann, 2004; Meinzer, Obleser, Flaisch, Eulitz, & Rockstroh, 2007). Our finding about the greatest contribution of the left middle frontal areas in logo-graphene representation for classification is also meaningful for clinical applications, such as intraoperative localization. Recently, precision estimation of brain networks revealed the phenomenon that archetype distributed network fractionate into multiple specialized networks in association regions which might inherit the accumulation of evolution (Braga, DiNicola, Becker, & Buckner, 2020; DiNicola & Buckner, 2021). An expansion-fractionation-specialization hypothesis has been consequently proposed, which are also supported by our findings here.

For logo-grapheme processing, brain activity in response to Chinese words and English words reading showed shared regions in the right superior temporal gyrus (rSTG), left middle frontal gyrus, right fusiform gyrus and rIFG. One interesting point should be noted that spatially very close but not overlapping clusters were found in the rSTG across any pairs of writing systems (Chi-W ⋂ Eng-W, rSTG, [60, −20, −10]; Chi-W ⋂ Pin, rSTG, [70, −16, −8]; Pin ⋂ Eng-W, rSTG, [66, −26, −6]). Previous studies demonstrated that the bilateral STG was involved in visual-audio integration both in English and Chinese reading (Holloway, van Atteveldt, Blomert, & Ansari, 2015; Kast, Bezzola, Jancke, & Meyer, 2011; McNorgan, Randazzo-Wagner, & Booth, 2013; Xia, 2020). Subtle anatomical separations supporting similar functions suggested that discrepant visual-audio linguistic features were integrated in subregions of the rSTG and supported the accommodation hypothesis. The overlap in brain activity located in the right fusiform gyrus might be considered a function of processing general features of objects or, more precisely, an adaptation of the second language, i.e., English, to the first language, i.e., Chinese. Notably, although previous studies considered the left ventral occipital-temporal cortex/fusiform gyrus to be universally involved in visual word form processing and was script invariant in monolingual individuals (Krafnick et al., 2016; Price, 2012), Gao and her colleagues found that subregions of the left ventral occipital-temporal cortex involved in visual word processing were different in Chinese-English bilingual individuals (Gao et al., 2017). In addition, researchers found that activation of the fusiform gyrus in English native speakers during reading was more left lateralized, but after learning Chinese, these individuals exhibited more bilateral fusiform activity during English reading (Mei et al., 2015). In this sense, the neural basis underlying Chinese as first language would affect brain responses to reading in the second language (Tan, 2005).

For brain processing of phonology, although nonoverlap was found between the brain activity during Chinese words and Chinese pinyin reading, shared brain activity for logo-grapheme were found in the rSTG, rIFG, right inferior parietal gyrus and right precentral gyrus, which are involved in visual-audio integration (Liu et al., 2006; Xia, 2020). Our results indicated that although phonological activations are mandatory, successful semantic access during Chinese reading depends more on morphological processing (C. S. Chen et al., 2009; Ipa et al., 2019; Ruan et al., 2018; Tan et al., 2005).

For the brain load of semantic processing, spatially closed clusters were found in the left inferior frontal gyrus for Chi-W ⋂ Pin (−38, 40, 6) and Pin ⋂ Eng-W (−38, 42, 10), which was in charge of general semantic processing (Y. Dong et al., 2005; Fiez, 1997; Petersen, Fox, Snyder, & Raichle, 1990; Tan et al., 2000). The overlapping in brain activity between Chinese words and Chinese pinyin reading for semantic processing was also found in the left middle temporal gyrus and left angular gyrus, which played the core role of ideographic reading. Pertinently, overlapping of semantic processing during Chinese pinyin and English words reading was shown in the left supramarginal gyrus, which proved important for alphabetic-related reading.

In addition to the neuroanatomical separations for different cognitive processing, we found significant interactions between language components and language types. Specifically, a balanced brain activity pattern was observed for Chinese word reading. Given that cue effects of phonetic radicals on pronunciation were not obvious in Chinese word reading (Perfetti, Zhang, & Berent, 1992; Zhou, 1978), the brain activity in response to phonological processing that was at a similar level to logo-grapheme and semantics proved that phonological codes in Chinese word reading are activated obligatorily, in line with previous studies (Perfetti & Zhang, 1995; Tan et al., 2005; Tan & Perfetti, 1999; Wu, Ho, & Chen, 2012). Interestingly, most activity in response to phonological representation was found in English words reading and the least activity was in response to Chinese pinyin reading. As alphabetic writing systems, both English word and Chinese pinyin reading require readers to access an exact sound by sequentially mapping letters/letter combinations onto phonemes and assembling them together (Ruan et al., 2018). However, Chinese pinyin (corresponding to two-character words in the current study) can be mapped directly to two monosyllables with tones, while English words can be mapped onto polysyllables with 5 to 9 phonemes. Furthermore, regarding orthographic consistency, Chinese pinyin, as a phonetic coding system, is highly transparent, but English words are relatively opaque (Y. P. Chen, Fu, Iversen, Smith, & Matthews, 2002; Fu, Chen, Smith, Iversen, & Matthews, 2002; Li et al., 2020; Perfetti & Dunlap, 2007; Ziegler et al., 2010). Thus, it is reasonable that during spelling, English words have a larger ‘lexical competition cohort’ and require more engagement for phonological processing, while Chinese pinyin completes grapheme-to-phoneme correspondence effortlessly (Y. P. Chen et al., 2002). It should be noted that the largest brain activity during semantic processing was found in Chinese pinyin reading, while the smallest was found in English word reading. For English words, once spelling was completed, exact sounds corresponded to unique semantics, utilizing the grapheme-phonology-semantics pathway (Plaut, McClelland, Seidenberg, & Patterson, 1996; Seidenberg & Mcclelland, 1989). Given the richness of homophones in Chinese (Kuo & Anderson, 2006; Packard, 2000), more resources would be needed for phonology-to-semantics mapping during Chinese pinyin word reading. In addition, based on the direct visual-orthography to semantics hypothesis for Chinese characters, Chinese pinyin processing might include both the grapheme (Chinese pinyin)-phonology-semantics pathway and the grapheme (Chinese pinyin)-phonology-logo-grapheme (Chinese word corresponded to Chinese pinyin)-semantics pathway (L. Chen, Perfetti, Fang, Chang, & Fraundorf, 2019; L. Chen, Perfetti, & Leng, 2019).

Previous brain imaging studies utilizing traditional univariate analyses failed to link neural responses to detailed language components and could only draw results at the cluster level. To the best of our knowledge, only one study conducted by Xu et al., used searchlight multivoxel pattern analysis to investigate voxel-wise neural response patterns of Chinese and English words reading in bilingual individuals (Xu et al., 2017). They found distributed neural populations in the left lateral occipital cortex, left fusiform gyrus, left temporal cortex, left temporal parietal cortex, left prefrontal cortex and superior parietal cortex serving for dominant language and the second language respectively, which to some extent supports our results. On the other side, with the development of deep learning and artificial intelligence, neural decoding techniques have achieved significant breakthroughs focused on single modality, such as visual information (Yoshida & Ohki, 2020) and articulatory movement (Anumanchipalli, Chartier, & Chang, 2019); however, there is still a large gap in decoding higher cognitive functions, such as language processing and decision making. Our findings shed light on the possibility of neural decoding for language comprehension.

Two limitations of this study should be addressed. Firstly, although searchlight RSA is a more fine-grained analysis method to depict brain responses, it can only determine linguistic feature-related processes but fails to indicate what manipulation has occurred. New methods or combining searchlight RSA with subtle experimental designs should be developed in future to reveal more details. Secondly, other factors, such as age at acquisition and language proficiency have been revealed impacting the brain responses during language processing, especially for bilinguals (Butler, 2012). Future bilingual studies should take those into consideration.

In summary, we found that the brain activity pattern of language-processing components in different writing systems showed neuro-anatomically distinct spatial distributions. Brain activity patterns were significantly correlated with reaction time and could reversely predict language types. Meanwhile, typical language-related regions had similar brain activation patterns and cognitive component processing of the three language types. Taken together, both separated and shared brain activity patterns were associated with linguistic features of each language system. Our findings supported and enriched the assimilation-accommodation hypothesis and demonstrated brain adaptation to long-term and complex language practice in bilingual individuals.

## Methods

### Participants

Fifty-one subjects (25 males; mean age = 23.4 years,) enrolled in the current study through online advertising. All were Chinese native speakers, had vision/corrected vision over 4.8, and met the inclusion criterion of no history of neurological disease or psychiatric disorders. All subjects underwent an 8-minute structural MRI scan and a task functional MRI scan that lasted approximately 40 minutes. The Edinburgh Handedness Inventory was used to identify participants’ handedness (Oldfield, 1971). Forty-one of all subjects were classified as right-handed, and 10 subjects had balanced dexterity. This study was approved by the Ethics Committee of the School of Life Sciences of Fudan University. Written informed consent was signed by every subject before the experiment.

### Stimuli and Task-fMRI Procedures

In current task-fMRI scans, an event-related design was utilized. Stimuli sets consisted of 3 conditions, Chinese word, Chinese pinyin, and English words, with 40 trials in each category. The Chinese word condition contain 40 two-character words; Chinese pinyin’s corresponding Chinese words were also two-character words. All stimuli were white and presented on black screen with horizontal visual angle as 4.37°. To prevent confounding effects, picture size, percentage of pixel, number of strokes, and word frequency were matched across conditions. There was no semantic equivalent in any two stimuli across conditions. All stimuli were visually presented for 1000 ms in a randomized order, and a fixation cross was presented for an interval of 4-6 s. Each stimulus subtended a visual angle of approximately 1 vertically and was presented in Song font for Chinese words and Arial for English words and Chinese pinyin in white against a black background. Participants were asked to respond to stimuli categories by pressing 3 different buttons with their index fingers. Notably, the current task did not require participants to work as fast as possible. Participants were asked to perform a semantic access task before a behavioral response task. For the semantic access task, subjects were asked to complete a recognition questionnaire after scanning. The subjects needed to identify whether the words in the checklist were shown in the fMRI task. The checklist comprises 120 two-character Chinese words containing 60 new words, 20 words from the Chinese word stimuli set, 20 words corresponding to the Chinese pinyin stimuli set and 20 words corresponding to the English word stimuli set. Given that memory performance was not the goal of the current study, the accuracy of the checklist was not used for further analysis. A practice experiment composed of 12 trials (4 stimuli of each condition) was performed before the normal fMRI scan was conducted to ensure that the subjects fully understood the tasks. Because of the machine fault, only 44 subjects’ button pressing performance was successfully recorded and used for subsequent behavioral analysis.

### Image Acquisition and Data preprocessing

Functional and anatomical images were acquired through a 3T Siemens Prisma scanner. An echo planar imaging (EPI) sequence was used for functional imaging data collection (echo time (TE) = 33 ms, flip angle =52°, matrix size = 110*96, field of view = 220*196 mm, slice thickness = 2 mm, number of slices = 72, and repetition time (TR) = 720 ms). For anatomical reference, a high-resolution T1-weighted image was acquired before tasks (TE = 2.56 ms, flip angle = 8°, matrix size = 320*320, field of view = 256*256 mm, slice thickness = 0.8 mm, number of slices =208, and TR =3000 ms).

Preprocessing and statistical analysis of fMRI data were performed using SPM12 (http://www.fil.ion.ucl.ac.uk/spm). For details, slice timing was first conducted to obtain temporal realignment with the middle EPI volume. Unwrapped spatial realignment was performed to correct nonlinear distortions from head movement and magnetic field inhomogeneity. Next, the T1 image was coregistered to the mean EPI image. The coregistered image was segmented and normalized to Montreal Neurological Institute (MNI) space to obtain deformation field parameters. All realigned EPI volumes were spatially normalized to MNI space by applying the deformation field parameters. Finally, spatial smoothing with a 6 mm full-width half-maximum isotropic Gaussian kernel was performed for the normalized volumes.

### Activation Analysis

In the first-level analysis, a general linear model (GLM) was used for fixed-effect analysis of each participant for each condition. For the GLM model, the convolution of stimulus onset time and canonical hemodynamic response function served as independent variables; the blood oxygen level-dependent (BOLD) time series signals served as dependent variables with 6 realignment parameters as regressors. After high-pass filtering, contrasts of interest were obtained for each condition relative to fixation. In the second-level analysis, one-sample t-tests were performed to obtain an activation map of contrasts for each condition, FDR correction (*p* < 0.05), cluster size > 10.

### Representational Similarity Analysis

RSA has been widely used to evaluate congruent patterns across modalities within/between subjects (Deniz, Nunez-Elizalde, Huth, & Gallant, 2019; Fischer-Baum et al., 2017; Kim, Liu, Liu, & Cao, 2020; Wang et al., 2018; L. Zhao et al., 2017). Specifically, in the current study, RSA bridged brain activity patterns and behavioral measurements in response to task stimuli (Kriegeskorte et al., 2008). Brain activity patterns were depicted by constructing neural representational dissimilarity matrices (neural RDMs) that have n*n dimensions (n = 40). Behavioral measurements were operationally defined by 3 behavioral representational dissimilarity matrices (logo-grapheme RDMs, phonological RDMs, and semantic RDMs), which are described in detail below.

#### Behavioral RDMs

To reveal the underlying internal neural mechanism for visual features and orthographic information processing, we constructed logo-grapheme RDMs by calculating one minus the overlapping ratio of basic units between any two stimuli in each condition. For Chinese words, the basic unit was a logo-grapheme that could not be semantically divided; for Chinese pinyin, the basic unit was a single “letter” or symbol for tone; and for English words, the basic unit was a single letter. Patterns for phonetic features were built by constructing phonological RDMs, calculated as one minus the ratio of shared phonetic units. The basic phonetic units were initials or finals or tones (regardless of position) for Chinese words and pinyin and vowels or consonants (regardless of position) for English words. To construct the semantic RDMs, a skip-gram model for the word2vector algorithm (the soft package was implemented in Python Gensim) was used to obtain the continuous vector representations for each stimulus. The dissimilarity value between any two stimuli was calculated by one minus the cosine angle between feature vectors corresponding to the stimuli. For Chinese word and pinyin, wiki-Chinese corpus was used as input. Parameters were set as window size = 5, negative sample number = 5, dimension number = 300, learning rate = 0.025, and subsampling rate = 1e-4. For English word, wiki-English corpus was used as input; parameters were set the same as they were for the Chinese word and pinyin reading assessment. The correlation among logo-grapheme, phonological and semantic RDMs across three writing systems were shown in Table S2.

#### Neural RDMs and Searchlight RSA

A standard GLM in first-level analysis was built to obtain trial-specific beta estimates for each subject. The first-level GLM contained 120 regressors corresponding to 40 stimuli in Chinese words, 40 stimuli in Chinese pinyin, and 40 stimuli in English words, with 6 head motion parameters regressed as potential confounding factors. All regressors were convolved with the canonical hemodynamic response function (HRF) and high-pass filtered by 128 s. After the first-level analysis, voxelwise beta-value maps were obtained for 120 stimuli in each subject. In searchlight RSA, each voxel was extended into a self-centered spherical region of interest (ROI) with a 6-mm radius. One-level beta-values within each ROI for each condition were extracted and dissimilarity was calculated as one minus Spearman’s rho between any two stimuli. After making the searchlight-center voxel pass through the cortex, we obtained 3 n*n dimensional neural RDMs corresponding to 3 conditions in each voxel and each subject (n=51). To decrease the calculation pressure and discard irrelevant noise, a gray matter mask was used with a probability higher than 0.2 in the Tissue Probability Map (TPM) atlas in SPM12. Additionally, we predefined a language-related anatomical mask including 15 cortical regions that have been reported to be involved in language comprehension-related processing and their symmetric regions in the right hemisphere. More specifically, the mas was composed of the bilateral middle frontal (5#, 6#), inferior frontal (9#, 10#, 11#, 12#), precentral (1#, 2#), supplementary motor (15#, 16#), inferior parietal (65#, 66#), supramarginal (67#, 68#), angular (69#, 70#), superior temporal (85#, 86#), middle temporal (89#,90#), superior temporal pole (87#,88#), middle temporal pole (91#, 92#), fusiform (59#, 60#), superior occipital (53#, 54#), middle occipital (55#, 56#), inferior occipital (57#, 58#) areas through the Automated Anatomical Labeling 3 (AAL3) template (Rolls, Huang, Lin, Feng, & Joliot, 2020; Rolls, Joliot, & Tzourio-Mazoyer, 2015). After discarding voxels in which variations in the time series of BOLD signals were less than 1/8 of the mean values, the overlap of the gray matter mask and language mask was defined as the final mask used in the following analysis. For each component, partial Spearman’s rank correlation was calculated between neural RDMs in each voxel and one behavioral RDM (e.g., Logo-grapheme RDM) with two other behavioral RDMs (e.g., the phonology RDM and semantic RDM) controlled for each condition (Chinese word, English word, and Chinese pinyin). Fisher’s R to Z transformation was further applied to access the whole-brain Z-map for each language component in each condition in each subject.

The degree of brain activity for each language component, i.e., brain loads, was defined as the sum of Z values in voxels whose neural RDM was significantly positively correlated with the behavioral RDM (Table S3). For validation, brain activity was also calculated as the number of voxels significantly representing each language component (Table S4).

### Support Vector Regression

To explore the association/dissociation of different cognitive brain loads derived from different writing system inputs, support vector regression (SVR) was applied (the soft package was implemented in Python Scikit-learn, https://scikit-learn.org/stable/). For each subject, brain loads corresponding to 3 language-component representations in 30 ROIs were extracted as feature vectors for each condition. On the other hand, categories of language-component representation were transformed into positive integers (Chinese word = 1, English word = 2, and Chinese pinyin = 3) as real label values. Thus, each subject contributed 3 feature samples with a size of 90 dimensions. A leave-one-subject-out (three samples contributed by each subject) cross-validation scheme was adopted to train the SVR model and test whether it could predict feature vector-related label values. To avoid information leakage, for each fold (3 samples/subject), RSA results of remain 50 subjects were used to generate group-level brain loads mask for logo-grapheme, phonology and semantics respectively. Training set and test set of brain loads were calculated based on fold-specific group-level mask. Furthermore, a recursive feature elimination (RFE) scheme was adopted to reveal what features make the representation mechanism strikingly different across different writing systems. For standard evaluation, unified group-level mask and leave-one-sample-out validation was adopted before conducting RFE. More specifically, each feature was eliminated, and the remaining features were fitted to the SVR model. The predictive contribution of a certain feature was measured by the root of the mean squared error (rMSE) of the SVR model fitted by the remaining 89 features. The higher the rMSE was, the worse the predictive performance was, and the more important absent features were. All features were sorted by absence-incurred rMSE in descending order, and absence-incurred rMSE smaller than the total rMSE (N = 90) was initially eliminated. Next, features (N = 90) were removed one by one in ascending order of absence-incurred rMSE, and then the remaining features (N = 89, 88, 87…) were used to fit the SVR model; then, the model was evaluated by negative rMSE. As a function of the remaining feature numbers N, the knee point of negative rMSE revealed the best number of features for the successful SVR model.

### Statistical analysis

Two-way repeated ANOVA was conducted to test significant differences across conditions and language components in reaction time, accuracy rate, and brain loads, with Bonferroni correction in post hoc tests. Spearman’s rank correlation was utilized to depict the relationship among each RDM. A linear mixed-effects model was performed to determine an adjusted reaction time to exclude intra-subject effects. Then, partial correlation was conducted to reveal the relationship between the brain loads and fitted reaction time with Bonferroni correction. One-tailed one-sample t-tests (H_0_: μ > 0) were performed across subjects to identify the voxels that were significantly involved in certain aspects of language component processing for every condition at the group level. The threshold was set to p < 0.05, uncorrected, and cluster size > 10.

## Acknowledgments

This work was supported by the Natural Science Foundation of China [grant numbers 81901826, 61932008], the Natural Science Foundation of Shanghai [grant number 19ZR1405600, 20ZR1404900], the Shanghai Municipal Science and Technology Major Project [No.2018SHZDZX01], ZJLab and Shanghai Center for Brain Science and Brain-inspired Technology.

## Author Contributions

J.F., M.C. and C.B. contributed to the design of the work; S.G. and W.G. contributed to the acquisition of data, S.G. and M.C. contributed to the analysis of data; M.C., C.B., L.T., J.F, S.G. and W.G. contributed to the interpretation of data; S.G., M.C. and W.G. drafted the work; M.C., L.T., K.X., T.J., S.G. and W.Z. substantively revised the work.

---

**Table S1.**
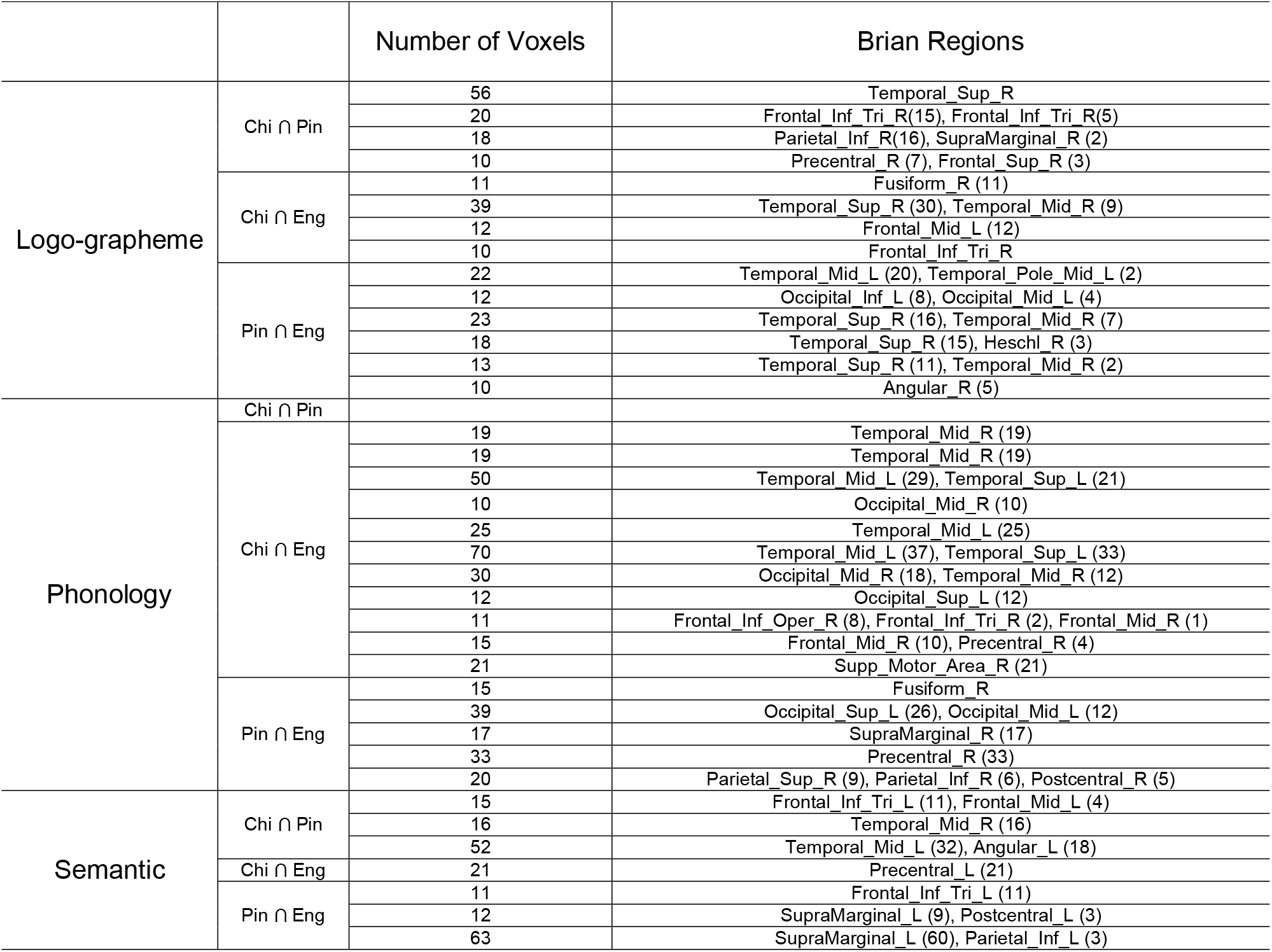
Clusters of information of overlap between any two conditions.

**Table S2.** Correlation among 3 components of RDMs across conditions.

|  | Chinese |  |  | English |  |  | Pinyin |  |  |
| --- | --- | --- | --- | --- | --- | --- | --- | --- | --- |
|  | Logo-graph<br>eme | Phonology | Semantic | Logo-graph<br>eme | Phonology | Semantic | Logo-graph<br>eme | Phonology | Semantic |
| Logo-graph<br>eme | - |  |  | - |  |  | - |  |  |
| Phonology | 0.048 | - |  | 0.371*** | - |  | 0.598*** | - |  |
| Semantic | -0.007 | 0.017 | - | 6.283e-04 | 0.047 | - | 0.109** | 0.109** | - |

\* indicates p<0.01; \*\*\* indicates p < 0.001, Bonferroni-corrected

**Table S3.** Brain activity elicited by each component in each ROI across conditions.

|  | Chinese |  |  | Pinyin |  |  | English |  |  |
| --- | --- | --- | --- | --- | --- | --- | --- | --- | --- |
|  | Logo-grapheme | Phonology | Semantic | Logo-grapheme | Phonology | Semantic | Logo-grapheme | Phonology | Semantic |
| Frontal_Mid_L | 1.34 | 0.61 | 5.37 | 3.34 | 0.71 | 18.73 | 9.7 | 7.91 | 1.02 |
| Frontal_Mid_R | 3.42 | 1.94 | 10.66 | 5.15 | 0.48 | 3.97 | 1.5 | 16.71 | 0.87 |
| Frontal_Inf_Oper_L | 0.18 | 1.27 | 1.63 | 0.23 | 0.14 | 2.75 | 0.14 | 1.51 | 0.55 |
| Frontal_Inf_Oper_R | 1.56 | 3.42 | 0.56 | 0.95 | 0.04 | 0.53 | 0.5 | 5.5 | 1.05 |
| Frontal_Inf_Tri_L | 0.41 | 3.15 | 1.97 | 3.77 | 0.06 | 12.86 | 2.8 | 8.22 | 2.07 |
| Frontal_Inf_Tri_R | 3.73 | 1.25 | 0.5 | 2.41 | 0.93 | 4.62 | 1.05 | 8.24 | 0.05 |
| Precentral_L | 3.22 | 2.92 | 1.39 | 0.64 | 0.97 | 6.95 | 3.44 | 13.84 | 2.58 |
| Precentral_R | 1.88 | 1.46 | 1.51 | 1.22 | 1.95 | 2.06 | 0.93 | 12.6 | 0.94 |
| Supp_Motor_L | 0.86 | 0.66 | 0.18 | 2.15 | 0.86 | 4.26 | 1.71 | 18.21 | 0.33 |
| Supp_Motor_R | 0.34 | 2.24 | 2.57 | 6.57 | 0.73 | 4.78 | 3.86 | 8.81 | 0.41 |
| Parietal_Inf_L | 1.03 | 0.12 | 1.06 | 3.61 | 0.31 | 6.54 | 2.32 | 8.94 | 0.97 |
| Parietal_Inf_R | 1.79 | 0.56 | 0.24 | 3.25 | 1.06 | 0.4 | 0.26 | 1.77 | 0.18 |
| SupraMarginal_L | 1.16 | 0.75 | 0.22 | 0.95 | 0.28 | 7.43 | 0.63 | 1.01 | 3.57 |
| SupraMarginal_R | 0.18 | 1.61 | 1.15 | 1.61 | 0.67 | 3.34 | 0.67 | 6.4 | 0.39 |
| Angular_L | 1.49 | 0.02 | 1.6 | 0.39 | 0.37 | 3.97 | 2.85 | 1.14 | 0.49 |
| Angular_R | 0.52 | 1.05 | 0.48 | 1.68 | 0.18 | 1.57 | 1.41 | 3.46 | 1.31 |
| Temporal_Sup_L | 1.5 | 3.82 | 2.57 | 0.97 | 0.58 | 2.84 | 2.16 | 7.86 | 0.88 |
| Temporal_Sup_R | 4.21 | 1.08 | 0.46 | 8.12 | 0.16 | 6.02 | 4.27 | 7.92 | 0.89 |
| Temporal_Pole_L | 2.4 | 1.31 | 0.52 | 1.6 | 0.58 | 0.89 | 3.59 | 1.88 | 0.63 |
| Temporal_Pole_R | 1.85 | 1.44 | 0.64 | 1.41 | 0.42 | 1.86 | 1.18 | 2.45 | 1 |
| Temporal_Mid_L | 3.63 | 9.95 | 3.89 | 5.14 | 1.01 | 22.53 | 9.21 | 16.89 | 2.83 |
| Temporal_Mid_R | 2.47 | 5 | 4.05 | 6.24 | 0.27 | 6.88 | 6.08 | 14.52 | 0.55 |
| Fusiform_L | 1.18 | 0.44 | 0.35 | 1.83 | 1.47 | 1.35 | 1.41 | 8.15 | 0.96 |
| Fusiform_R | 1.74 | 1.26 | 1.75 | 2.14 | 2.63 | 1.79 | 2.49 | 5.89 | 0.73 |
| Occipital_Sup_L | 0.2 | 0.43 | 0.86 | 2.23 | 1.78 | 2.4 | 0.13 | 4.33 | 0.12 |
| Occipital_Sup_R | 1.59 | 2.89 | 0.69 | 2.23 | 1.44 | 9.34 | 4.8 | 7.58 | 1.76 |
| Occipital_Mid_L | 0.35 | 1.23 | 0.49 | 1.83 | 0 | 0.4 | 2.36 | 3.27 | 0.56 |
| Occipital_Mid_R | 1.2 | 0.31 | 0 | 0.63 | 0.58 | 6.49 | 0.86 | 4.8 | 0.25 |
| Occipital_Inf_L | 1.53 | 1.39 | 0.36 | 1.66 | 0.08 | 3.02 | 0.75 | 14.75 | 6.32 |
| Occipital_Inf_R | 0.11 | 0 | 0.03 | 1 | 0 | 0.54 | 0.91 | 5.04 | 1.1 |

**Table S4.** Voxel numbers involved in each component in each ROI across conditions.

|  | Chinese |  |  | Pinyin |  |  | English |  |  |
| --- | --- | --- | --- | --- | --- | --- | --- | --- | --- |
|  | Logo-grapheme | Phonology | Semantic | Logo-grapheme | Phonology | Semantic | Logo-grapheme | Phonology | Semantic |
| Frontal_Mid_L | 41 | 19 | 150 | 88 | 18 | 460 | 280 | 213 | 29 |
| Frontal_Mid_R | 102 | 57 | 282 | 143 | 14 | 104 | 41 | 430 | 21 |
| Frontal_Inf_Oper_L | 6 | 37 | 45 | 6 | 4 | 65 | 4 | 39 | 16 |
| Frontal_Inf_Oper_R | 47 | 100 | 16 | 25 | 1 | 13 | 14 | 139 | 29 |
| Frontal_Inf_Tri_L | 11 | 94 | 55 | 92 | 2 | 303 | 78 | 210 | 55 |
| Frontal_Inf_Tri_R | 115 | 39 | 14 | 60 | 21 | 110 | 29 | 203 | 1 |
| Precentral_L | 90 | 80 | 39 | 15 | 23 | 154 | 90 | 328 | 69 |
| Precentral_R | 53 | 42 | 43 | 32 | 48 | 52 | 23 | 315 | 23 |
| Supp_Motor_L | 24 | 20 | 5 | 56 | 21 | 98 | 46 | 436 | 10 |
| Supp_Motor_R | 10 | 66 | 69 | 161 | 20 | 119 | 105 | 225 | 13 |
| Parietal_Inf_L | 30 | 4 | 30 | 86 | 8 | 157 | 64 | 224 | 28 |
| Parietal_Inf_R | 56 | 19 | 7 | 77 | 27 | 9 | 7 | 44 | 4 |
| SupraMarginal_L | 33 | 24 | 6 | 22 | 7 | 170 | 18 | 28 | 96 |
| SupraMarginal_R | 6 | 50 | 33 | 41 | 20 | 79 | 19 | 168 | 11 |
| Angular_L | 45 | 1 | 46 | 10 | 12 | 111 | 77 | 30 | 14 |
| Angular_R | 16 | 34 | 13 | 45 | 5 | 44 | 42 | 86 | 39 |
| Temporal_Sup_L | 47 | 115 | 66 | 25 | 16 | 68 | 62 | 209 | 24 |
| Temporal_Sup_R | 129 | 33 | 13 | 203 | 5 | 149 | 123 | 216 | 25 |
| Temporal_Pole_L | 74 | 40 | 15 | 43 | 14 | 22 | 104 | 52 | 17 |
| Temporal_Pole_R | 57 | 42 | 17 | 37 | 12 | 47 | 34 | 68 | 27 |
| Temporal_Mid_L | 102 | 295 | 112 | 129 | 28 | 544 | 250 | 438 | 75 |
| Temporal_Mid_R | 77 | 145 | 117 | 162 | 8 | 174 | 166 | 377 | 15 |
| Fusiform_L | 35 | 14 | 9 | 42 | 38 | 34 | 39 | 203 | 27 |
| Fusiform_R | 54 | 37 | 49 | 52 | 69 | 44 | 65 | 145 | 18 |
| Occipital_Sup_L | 6 | 12 | 23 | 50 | 41 | 53 | 5 | 105 | 3 |
| Occipital_Sup_R | 44 | 84 | 18 | 47 | 34 | 197 | 111 | 175 | 43 |
| Occipital_Mid_L | 9 | 34 | 11 | 40 | 0 | 9 | 60 | 73 | 13 |
| Occipital_Mid_R | 31 | 9 | 0 | 14 | 14 | 140 | 22 | 110 | 5 |
| Occipital_Inf_L | 38 | 42 | 10 | 39 | 2 | 66 | 20 | 342 | 151 |
| Occipital_Inf_R | 3 | 0 | 1 | 20 | 0 | 10 | 20 | 102 | 25 |

## Notes

### Competing Interest Statement

The authors have declared no competing interest.

